# Neutralization of B.1.1.28 P2 variant with sera of natural SARS-CoV-2 infection and recipients of BBV152 vaccine

**DOI:** 10.1101/2021.04.30.441559

**Authors:** Gajanan Sapkal, Pragya D. Yadav, Raches Ella, Priya Abraham, Deepak Y. Patil, Nivedita Gupta, Samiran Panda, V. Krishna Mohan, Balram Bhargava

## Abstract

The emergence of new SARS-CoV-2 variants has been a serious threat to the public health system and vaccination program. The variant of concerns have been the under investigation for their neutralizing potential against the currently available COVID-19 vaccines. Here, we have determined the neutralization efficacy of B.1.1.28.2 variant with the convalescent sera of individuals with natural infection and BBV152 vaccination. The two-dose vaccine regimen significantly boosted the IgG titer and neutralizing efficacy against both B.1.1.28.2 and D614G variants compared to that seen with natural infection. The study demonstrated 1.92 and 1.09 fold reductions in the neutralizing titer against B.1.1.28.2 variant in comparison with prototype D614G variant with sera of vaccine recipients and natural infection respectively.

## Text

India has reported cases infected with the SARS-CoV-2 UK variant (B.1.1.7).^1^ Recently, South Africa variant (B.1.351) and Brazil variant P2 lineage (B.1.1.28.2) have also been detected in international travellers travelling to India from abroad. The impact on the emergence of these new variants on the efficacy of the currently available COVID-19 vaccines or neutralizing capability of the sera of individuals infected naturally with the earlier circulating strains is currently under investigation. Although some of the vaccines seem to be effective against the UK variant,^1,2,3^ the efficacy of them against the South African variant has been demonstrated to be less efficacious.^2,4^ A SARS-CoV-2 vaccine that used inactivation platform has been reported to be 50.7% efficacious from Brazil, where the B.1.1.28.2 variant is moreprevalent (NCT0445659). In this study, we determined the IgG immune response and neutralizing activity of the 19 convalescent sera specimens obtained from the recovered cases of COVID-19 and confirmed for B.1.1.7, B.1.351, B.1.1.28.2 (n=2 each), B1 lineage (n=13) (15-113 days post positive test) and from 42 participants immunized with an inactivated Covid-19 vaccine, BBV 152 (Covaxin) as part of a phase II clinical trial (two months post the second dose).^5^ IgG response was determined using the spike protein (S1-RBD) and nucleocapsid (N) protein-based ELISAs. Neutralizing antibody (NAb) titers of all the serum specimens were evaluated against B.1.1.28.2 variant compared to prototype D614G variant using plaque reduction neutralization test (PRNT50) [Supplemental information].

The geometric mean titer (GMT) of an IgG titer for S1-RBD and N protein ELISA was observed to be 794.8 and 4627 respectively for the SARS-CoV-2 recovered individuals. Covaxin recipients showed a GMT IgG titer of 2250 with S1-RBD and 3099 with the N protein compared; the former being significantly high compared to natural infection (Figure 1A). The geometric mean titer (GMT) of the neutralizing antibodies (NAb) of the sera with natural infection and Covaxin recipients for the prototype D614G strain was 120.1 and 337.5. In the B.1.1.28.2 variant, the GMT for individuals with natural infection was observed to be 109.2, while that of vaccine recipient was found to be 175.7.

**Figure 1.**
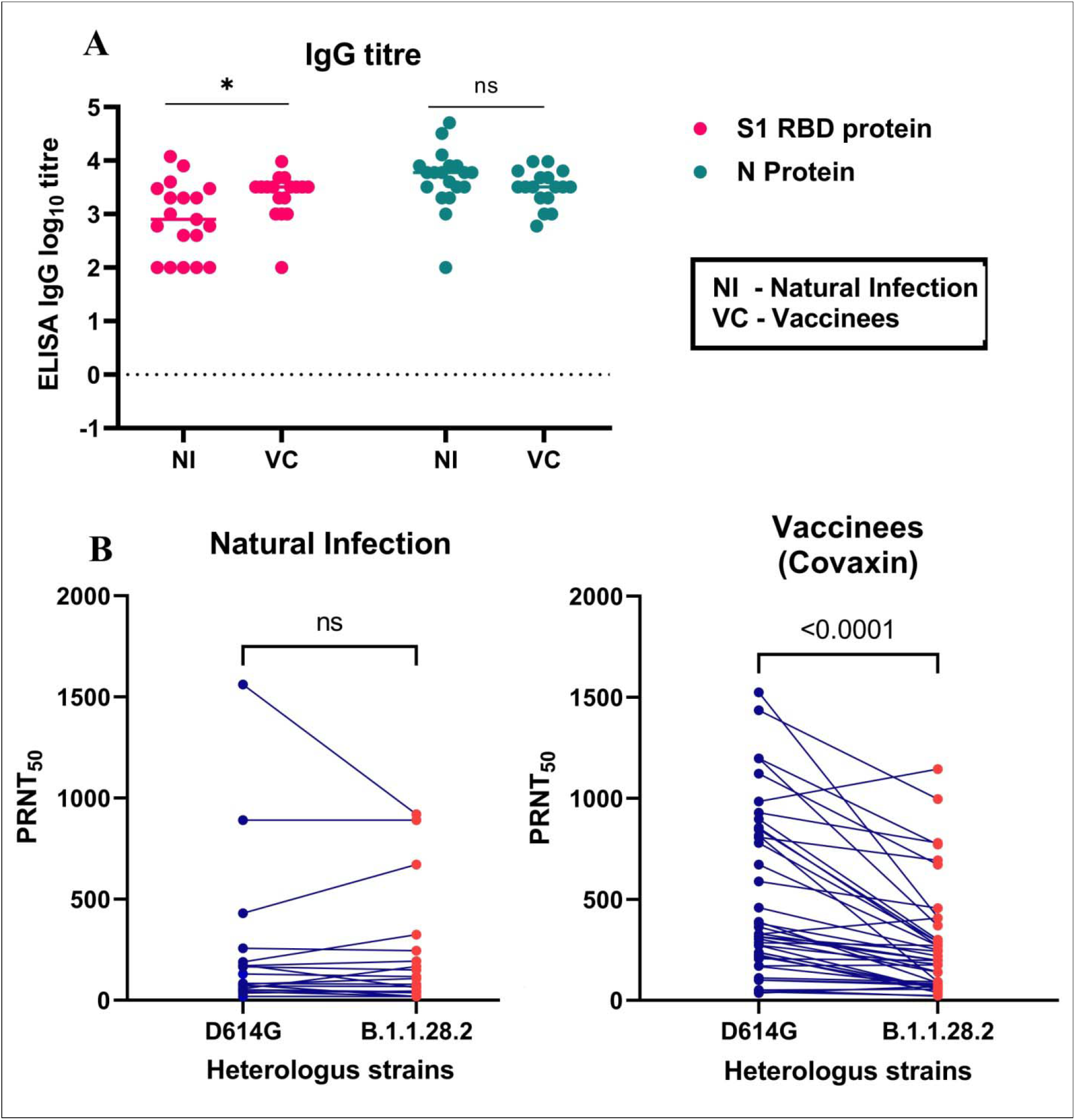
IgG response and neutralizing activity post-natural SARS-CoV-2 infection and Covaxin recipients. **Panel A** depicts the results of S1-RBD protein (Pink circles) and N protein (Blue circles) based SARS-CoV-2 IgG ELISA to determine the IgG titer in sera of naturally infected individuals (n=19) and Covaxin recipients (n=42). Mann-Whitney test was used for the comparison. **Panel B** shows the results of the plaque reduction neutralization test to assess neutralization activity of the prototype virus D614G (Blue circles) with B.1.1.28 P2 variant (Red circles) with sera of naturally infected individuals (n=19) and Covaxin recipients (n=42). The natural Infection had two samples each from Brazil, South Africa and the UK and thirteen of B1 lineage. Wilcoxon matched-pairs signed-rank test was used to compare the significance. A significant reduction was observed in the samples of the vaccines, whereas a non-significant change was observed in the cases with natural Infection.

This study shows that the two-dose Covaxin regimen significantly boosted the IgG titer and neutralizing efficacy against both the variants compared to that seen with natural infection. A two-tailed Wilcoxon paired signed-ranks test demonstrated a significant difference between prototype D614G strain and B.1.1.28.2 variant (Figure 1B). Results confirm 1.92 and 1.09 fold reductions in the neutralizing titer against B.1.1.28.2 variant in comparison with prototype D614G variant with sera of vaccine recipients and natural infection respectively.

The reduced neutralizing capability of vaccine’s sera has been reported earlier.^2^ Findings from another inactivated vaccine recently reported no neutralization to the P1 (B.1.1.28), albeit sera samples assessed were five months post the second dose.^6^ This study further corroborates recent findings indicating high levels of cross-reactivity in sera collected from variant infected individuals.^7^

## Ethical approval

The study is approved by the Institutional Biosafety Committee and Institutional Human Ethics Committee of ICMR-NIV, Pune, India

## Author Contributions

PDY, RE and PA contributed to study design, data collection, data analysis, interpretation and writing and critical review. GS and DYP contributed to data collection, interpretation, writing and critical review. NG, SP, VKM and BB contributed to critical review and finalization of the paper.

## Acknowledgement

We thank the scientific staff of ICMR-NIV, Pune Dr. Anita Shte, Dr. Rima Sahay, Dr. Gururaj Deshpande, Dr. Dimpal Nyayanit and Dr. Abhinendra Kumar for providing excellent support. Authors gratefully acknowledge the staff of ICMR-NIV, Pune including Mr. Prasad Sarkale, Ms. Pranita Gawande, Mrs. Ashwini Waghmare, Ms. Jyoti Yemul and Ms. Manisha Dudhmal for extending excellent technical support.

