## Supplemental information for "Neutralization of B.1.1.28 P2 variant with sera of natural SARS-CoV-2 infection and recipients of BBV152 vaccine"

### Detection of anti-SARS-CoV-2 IgG antibodies using Spike and nucleocapsid protein based-ELISA

Briefly, 96-well ELISA plates (Nunc, Germany) were coated with SARS-CoV-2 specific antigens (S1-RBD at a concentration of 1.5µg/well and N protein at a concentration of 0.5µg/well in PBS pH 7.4). The plates were blocked with a Liquid Plate Sealer (CANDOR Bioscience GmbH, Germany) and Stabilcoat (Surmodics) for two hours at 37°C. The plates were washed two times with 10 mM PBS, pH 7.4 with 0.1 per cent Tween-20 (PBST) (Sigma-Aldrich, USA). Starting at a 1:50 dilution sera of the international travelers, SARS-CoV-2 infected individuals and the Covaxin recipients were serially four-fold diluted and added to the antigen-coated plates and incubated at 37°C for one hour. These wells were washed five times using 1× PBST and followed by addition 50 μl/well of anti-human IgG horseradish peroxidase (HRP) (Sigma) diluted in Stabilzyme Noble (Surmodics). The plates were incubated for half an hour at 37°C and then washed as described above. Further, 100 μl of TMB substrate was added and incubated for 10 min. The reaction was stopped by 1 N H_2_SO_4_, and the absorbance values were measured at 450 nm using an ELISA reader. Anti-SARS-CoV-2 antibody (NIBSC code 20/130) was also included in the assay as a reference standard. The cut-off for the assays was set at twice of average OD value of negative control. The endpoint titer of a sample is defined as the reciprocal of the highest dilution that has a reading above the cutoff value.

### Detection of neutralizing antibodies using plaque reduction neutralization test

A four-fold serial dilution of the serum samples were mixed with an equal amount of each virus suspension (50-60 Plaque forming units in 0.1 ml) separately. After incubating the mixtures at 37°C for 1 hr, each virus-diluted serum sample (0.1 ml) was inoculated onto duplicate wells of a 24-well tissue culture plate containing a confluent monolayer of Vero CCL-81 cells. After incubating the plate at 37°C for 60 min, an overlay medium consisting of 2% Carboxymethyl cellulose (CMC) with 2% fetal calf serum (FCS) in 2× MEM was added to the cell monolayer, and the plate was further incubated at 37°C in 5% CO_2_ for 5 days. Plates were stained with 1% amido black for an hour. Antibody titers were defined as the highest serum dilution that resulted in >50 (PRNT_50_) reduction in the number of plaques.
